# Cell type- and state- resolved immune transcriptomic profiling identifies glucocorticoid-responsive molecular defects in multiple sclerosis T cells

**DOI:** 10.1101/2022.06.29.498195

**Authors:** Tina Roostaei, Afsana Sabrin, Pia Kivisäkk, Cristin McCabe, Parham Nejad, Daniel Felsky, Hanane Touil, Ioannis S. Vlachos, Daniel Hui, Jennifer Fransson, Nikolaos A. Patsopoulos, Vijay K. Kuchroo, Violetta Zujovic, Howard L. Weiner, Hans-Ulrich Klein, Philip L. De Jager

## Abstract

The polygenic and multi-cellular nature of multiple sclerosis (MS) immunopathology necessitates cell-type-specific molecular studies in order to improve our understanding of the diverse mechanisms underlying immune cell dysfunction in MS. Here, by generating a dataset of 1,075 transcriptomes from 209 participants (167 MS and 42 healthy), we assessed MS-associated transcriptional changes in six implicated cell-type-states: naïve and memory helper T cells and classical monocytes purified from peripheral blood, each in their primary (*ex vivo*, unstimulated) and *in vitro* stimulated states. Our data suggest that primary profiles show larger MS-associated differences than the post-stimulation contexts. We further identified shared and distinct changes in individual genes, biological pathways, and co-expressed gene modules in MS T cells and monocytes, and prioritized genes such as *ZBTB16* as MS-associated regulators in both cell types. Of six identified MS-associated co-expressed gene modules, three (two lymphoid and one myeloid) were replicated in independent data from peripheral blood mononuclear cells (PBMC) and monocyte-derived macrophages. A subsequent *in silico* drug screen prioritized small-molecule compounds for reversing the perturbation of the MS-associated modules. The effects of glucocorticoid receptor agonists as the top-identified therapeutic class for the replicated T cell modules were validated using targeted *in silico* analyses and *in vitro* experiments, suggesting the coordinated dysregulation of glucocorticoid-responsive genes in MS T cells. In summary, our study identifies and validates individual genes and co-expressed gene modules from T and myeloid cells that are perturbed in MS, offering new targets for therapeutic discovery and biomarker development to guide the management of MS.

## Introduction

Multiple sclerosis (MS) is a chronic inflammatory disease of the central nervous system (CNS). The earlier phase of the disease is characterized by episodes of focal blood-brain-barrier breakdown and recruitment of peripheral lymphoid and myeloid cells to active demyelinating areas within the white and gray matter, in addition to slow accumulation of lymphocytes in the meninges leading to further inflammation and demyelination in the underlying cortex^1^. The enrichment for MS susceptibility genes among most lymphoid and myeloid cell subtypes further strengthens the evidence for several different peripheral immune cell types to play a role in MS pathogenesis^2^.

The main therapeutic mechanisms currently applied to MS treatment include modulating the function of immune cells, depleting subtypes of these cells, or retaining them in the periphery and preventing them from entering the CNS^3^. However, none of these interventions provide definitive treatment, highlighting our limited understanding of the molecular defects in immune cells that lead to MS. Previous transcriptomic studies on whole blood or peripheral blood mononuclear cells (PBMC) have prioritized a number of genes and pathways in relation to immune cell dysfunction in MS^4–6^. However, the inference of cell-type-specific defects from these samples that contain variable proportions of different cell types remains a suboptimal approach.

More recently, data collection from purified peripheral immune cell types has started to address this problem and to improve our understanding of the cell-type-specific immune dysfunction associated with MS^7–10^. Here, overcoming some of the limitations of prior studies (including limited sample sizes, treatment-related confounding effects, and limited profiling of more than one cell type/state at a time), we aimed to disentangle shared and distinct gene expression changes (both at the individual gene level and gene networks) in 6 immune cell types and states implicated in MS. By generating and analyzing a large dataset of 1,075 transcriptomes from 209 individuals, we assessed MS-associated transcriptional changes in three cell types: naïve and memory helper T cells and classical monocytes, each in two states: primary (*ex vivo*) and stimulated. We then complemented our analyses with replication studies, *in silico* drug screen, and *in vitro* validation experiments. Our results, (1) provide cell-type-specific information on genes and pathways perturbed in MS, (2) identify and replicate coordinated gene expression changes associated with MS, and (3) identify and validate a connection between coordinated transcriptional changes in MS T cells and the therapeutic effect of glucocorticoids.

## Results

### Data generation

MS participants were selected from the Comprehensive Longitudinal Investigation of Multiple Sclerosis at the Brigham and Women’s Hospital (CLIMB) study^11^ using the following selection criteria: (1) age 18-60 years old, (2) a diagnosis of MS fulfilling 2010 McDonald criteria, (3) a relapsing-remitting (RR) disease course at the time of sampling, (4) being untreated or on treatment with glatiramer acetate (GA) for >6 months at the time of sampling, (5) no evidence of disease activity in the prior 6 months, (6) no steroid use in the preceding 30 days, and (7) an Expanded Disability Status Scale (EDSS) score between 0 to 3. Of note, we restricted the inclusion of treated MS individuals to those receiving GA in order to minimize the confounding effects of treatment on immune cell gene expression, as we have previously demonstrated a lack of differential gene expression in PBMC of GA-treated and untreated MS individuals^12^. Healthy participants were selected from the PhenoGenetic Project, a resource of individuals free of inflammatory or infectious diseases that have previously been used to study the extent of variation in immune function among healthy individuals in the ImmVar study^13,14^.

A prospectively collected, cryopreserved vial of PBMC was accessed for each participant. PBMC were thawed and sorted using magnetic beads followed by flow cytometry (see Methods), and three cell populations were retained: CD14^+^CD16^+^ (classical) monocytes as well as naïve and memory CD4^+^ (helper) T cells. Each of the three cellular samples were then split into two aliquots, with one aliquot remaining unstimulated while the other aliquot was placed in culture and stimulated before RNA was extracted for sequencing. Anti-CD3/CD28 antibodies were used for T cell stimulation in order to mimic stimulation by antigen-presenting cells^15^, and TNF-α was used for monocyte activation given its implication in MS pathophysiology^16^. Thus, 6 RNA samples (3 cell types, profiled unstimulated and stimulated) were collected from each cryopreserved PBMC sample (Figure 1). All RNA samples were processed using the Smart-Seq2 protocol to produce cDNA and a transcriptomic profile using sequencing. A rigorous data preprocessing pipeline was instituted after the experimental pipeline was concluded (see Methods), and, ultimately, data were retained from 209 participants of European ancestry. These include 42 healthy participants as well as 167 individuals with RRMS. The latter included 69 untreated and 98 GA-treated participants. The demographic characteristics of the participants are presented in Table 1.

**Figure 1.**
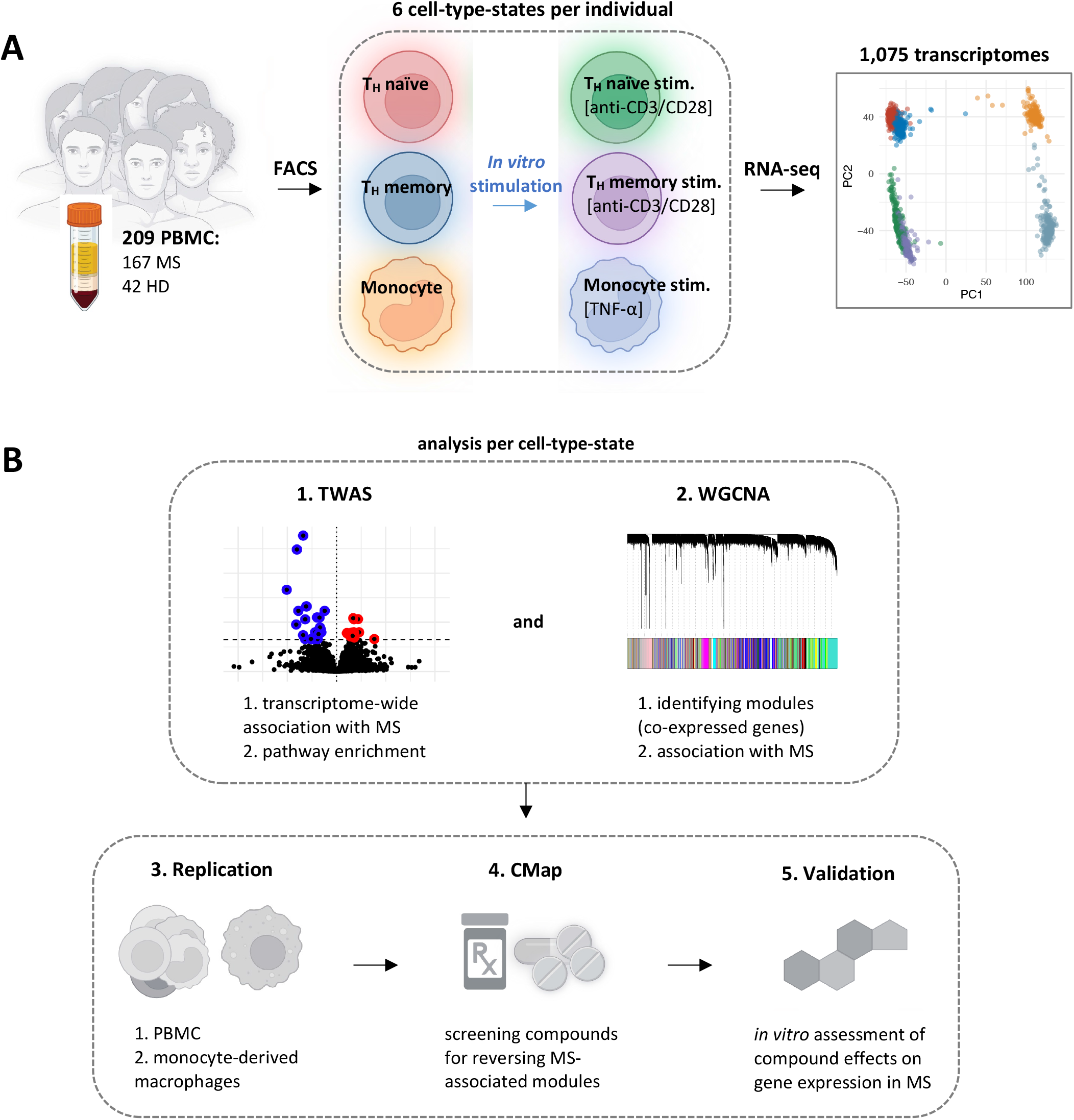
Summary of the generated data and the analyses performed in the study. (A) After quality control, 1,075 transcriptomes generated from 209 individuals and 6 cell-type-states were retained for analysis. (B) Analysis pipeline included identification of MS-associated genes, pathways, and co-expressed gene modules in each cell-type-state, followed by replication studies, *in silico* drug screen, and *in vitro* validation experiments.

**Table 1.** Demographic characteristics of the participants.

| study | cell type | diagnosis | MS medication | MS age | MS sex | HD age | HD sex |
| --- | --- | --- | --- | --- | --- | --- | --- |
| main study | six cell-type-states | 167 MS, 42 HD | 69 untreated, 98 GA | 40 ± 7 | 124F, 43M | 37 ± 9 | 31F, 11M |
| replication #1 | PBMC | 29 MS, 36 HD | 29 GA | 40 ± 10 | 17F, 12M | 35 ± 9 | 26F, 10M |
| replication #2 | monocyte-derived macrophages | 28 MS, 11 HD | 11 untreated, 17 treated* | 38 ± 11 | 24F, 4M | 37 ± 11 | 10F, 1M |
| *in vitro* experiment | PBMC | 5 MS | 2 untreated, 3 treated** | 52 ± 16 | 3F, 2M | N/A | N/A |

Age is reported in mean (years old) ± standard deviation.
\*treatments: n=4 dimethyl fumarate, n=3 natalizumab, n=3 fingolimod, n=2 rituximab, n=2 interferon-beta, n=1 teriflunomide, n=1 methotrexate, n=1 glatiramer acetate.
\*\*treatments: n=1 teriflunomide, n=2 glatiramer acetate.
F: female; GA: glatiramer acetate; HD: healthy donor; M: male; MS: multiple sclerosis; PBMC: peripheral blood mononuclear cells.

### The effects of diagnosis and treatment on T cell and monocyte gene expression in MS

For each of the 6 cell-type-states, we first performed transcriptome-wide differential gene expression analysis between (1) untreated MS and healthy participants, (2) GA-treated MS and healthy participants, and (3) untreated and GA-treated MS participants. The number of significant differentially expressed genes (FDR-adjusted *p* <0.05) in each comparison is shown in Figure 2A. Two main conclusions can be drawn from these simple, descriptive analyses:

1. There is little or no significant difference in gene expression (range: 0-7 significant genes) between MS (either untreated or GA-treated) and healthy participants in stimulated T cells and monocytes. On the other hand, unstimulated cell types showed higher number of differentially expressed genes. Interestingly, memory T cells (Tmem), which are produced from prior *in vivo* activation of naïve T cells (Tn), had lower number of differentially expressed genes in comparison to their naïve counterparts.
2. We did not observe notable difference in gene expression in any of the studied primary (unstimulated) and stimulated cell types between untreated and GA-treated MS participants (range: 0-5 significant genes; Supplementary Table 1). This observation was further supported by the high correlations between *t*-statistics from the transcriptome-wide results of the comparisons between untreated MS vs. healthy and GA-treated MS vs. healthy participants (Figure 2B, Supplementary Figure 1). These results corroborate previous literature on the lack of transcriptional difference between untreated and GA-treated MS in PBMC^12^. As a result of these observations, we decided to combine data from untreated and GA-treated MS participants into one group (referred to as the MS group), and take advantage of the larger sample size for performing gene expression comparisons between MS and healthy participants.

**Figure 2.**
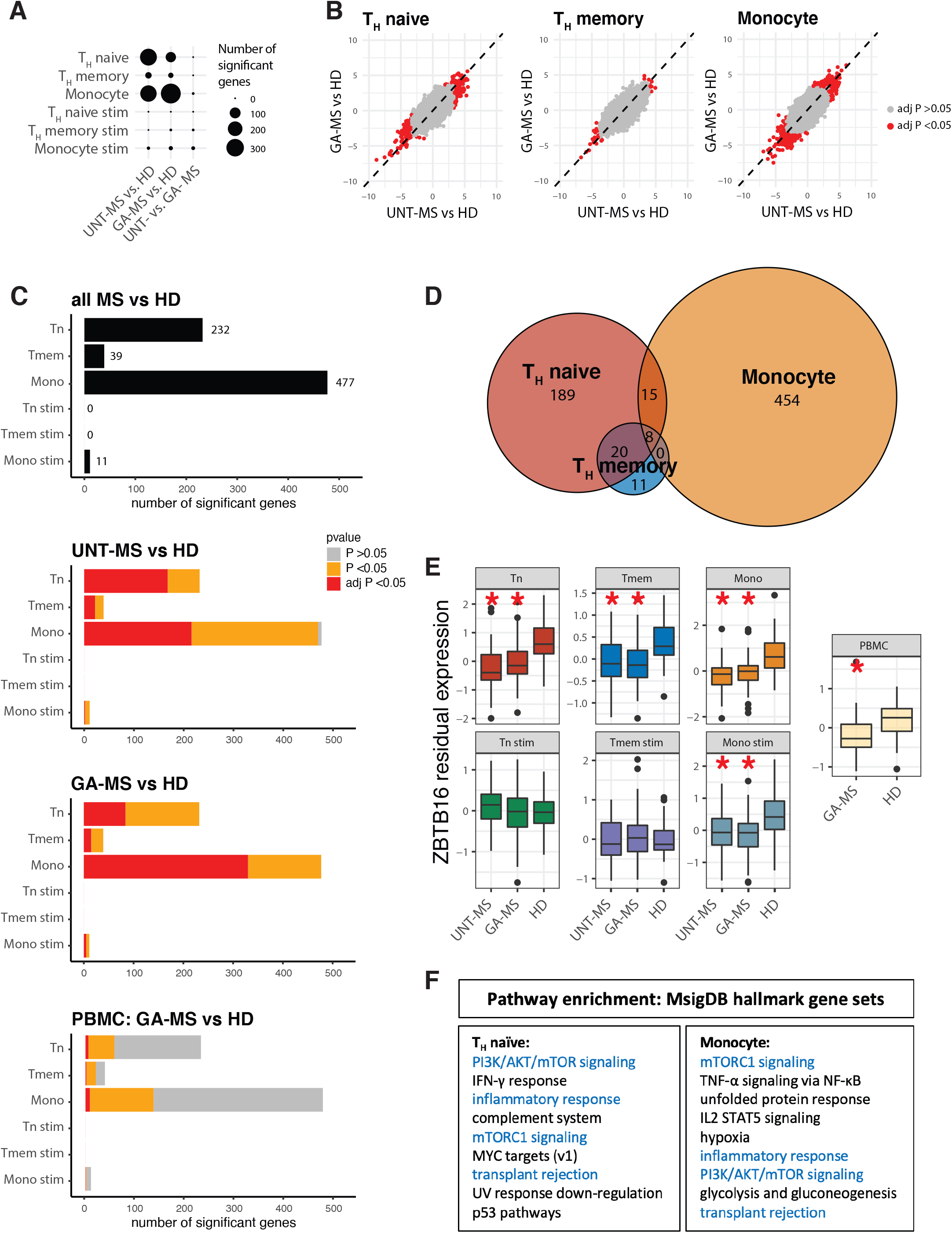
Results from transcriptome-wide association studies. (A) Number of significant genes (FDR-adjusted *p* <0.05) in transcriptome-wide association studies comparing each pair of the 3 participant groups (untreated MS, GA-treated MS, and healthy individuals) in each of the 6 studied cell-type-states. (B) Correlations between *t*-statistics from the transcriptome-wide association studies comparing the 2 MS groups with healthy individuals. Significant genes in any of the associations are shown in red. (C) Number of significant genes associated with MS in each cell-type-state are shown using black bars. Significance of the associations between each of these genes and MS in comparisons performed between each MS group and healthy participants, and also in comparisons between MS and healthy PBMC samples are shown in colors: FDR-adjusted *p* <0.05 with consistent direction of effect is shown in red, nominal *p* <0.05 with consistent direction of effect is shown in orange, and others are shown in gray. (D) Number of shared and distinct genes associated with MS in different cell types. (E) Expression of *ZBTB16* in each of the 6 cell-type-states, in addition to PBMC. Red stars represent FDR-adjusted *p* <0.05 for comparisons between MS and healthy participants. (F) Pathways enriched in MS-associated genes in T cells and monocytes. Shared pathways are shown in blue. UNT-MS: untreated MS, GA-MS: GA-treated MS, HD: healthy donor.

### T cell and monocyte genes differentially expressed in MS

Performing transcriptome-wide differential gene expression analysis between MS (combined group) and healthy participants, we identified 704 genes that were differentially expressed (FDR-adjusted *p* <0.05) in at least one of the 6 cell-type-states (Figure 2C, Supplementary Table 2, Supplementary Data). The vast majority of the identified differentially expressed genes (>99%) had a nominal *p* <0.05 with a consistent direction of effect in both untreated MS vs. healthy and GA-treated MS vs. healthy comparisons in their relevant cell-type-states. However, only 50% of the genes had an FDR-adjusted *p* <0.05 in each of the two separate comparisons (Figure 2C), highlighting the effect of the larger sample size in identifying larger number of differentially expressed genes.

We then performed differential gene expression analysis on RNA-seq data available from a smaller number of PBMC samples (n=65; 29 GA-treated MS and 36 healthy participants, from which, 6 of the MS and 29 of the healthy individuals had paired data in the 6 cell-type-states dataset; details in the Methods, Table 1). Of the 704 identified genes, we replicated the association of 12 genes (FDR-adjusted *p* <0.05 with consistent direction of effect), and 170 additional genes had a nominal *p* <0.05 in the PBMC samples (Figure 2C, Supplementary Table 2), suggesting a broader replication of our results in this small replication dataset. However, the lower correlations between *t*-statistics from the PBMC analysis and those from the purified cell types – which are part of the PBMC mixture – suggests that the differences observed in primary cell types in MS cannot be fully captured using PBMC data, and vice versa (Supplementary Figure 2).

Another observation from our data was that although a large percentage (72%) of the Tmem differentially expressed genes were also differentially expressed in Tn cells, this was not the case between Tn cells and monocytes (Figure 2D). Of the >200 differentially expressed genes in MS identified in Tn cells and >400 differentially expressed genes in monocytes, only 23 genes were shared between these two cell types (Table 2). Among these genes, *ZBTB16* and *HMGB2* were differentially expressed in all 3 primary cell types, in addition to PBMC. *ZBTB16* was also differentially expressed in stimulated monocytes (Figure 2E). *ZBTB16* (Zinc finger and BTB domain-containing 16), also known as *PLZF* (Promyelocytic Leukemia Zinc Finger) or *ZNF145*, has previously been reported to be a marker associated with MS and a number of other autoimmune diseases: hypomethylation in a *ZBTB16* enhancer region is reported in PBMC in MS^17^, and *ZBTB16* downregulation is observed in systemic lupus erythematosus (SLE) B cells^18^ and type 1 diabetes (T1D) PBMC^19^. Our results on the downregulation of *ZBTB16* in T cells, monocytes, and PBMC in MS are in line with the previous literature on this gene and suggest an important role for *ZBTB16* in dysregulated immune response leading to autoimmunity.

**Table 2.** Genes associated with MS in both T cells and monocytes.

| HGNC symbol | Ensembl ID | chromosome | t stats. Tn | t stats. Tmem | t stats. Mono | t stats. PBMC |
| --- | --- | --- | --- | --- | --- | --- |
| ACTR3 | ENSG00000115091 | 2 | **3.74** | 3.2 | **5.95** | 2.84 |
| BRD7 | ENSG00000166164 | 16 | **3.46** | -0.02 | **4.87** | 0.94 |
| CASS4 | ENSG00000087589 | 20 | **-4.74** | **-4.89** | **-3.48** | -2.56 |
| CCND3 | ENSG00000112576 | 6 | **-6.35** | **-4.69** | **-5.01** | -3.08 |
| CIRBP | ENSG00000099622 | 19 | **-3.48** | -1.9 | **-4.65** | -2.29 |
| DDIT4 | ENSG00000168209 | 10 | **-5.53** | -2.93 | **-5.73** | -3.83 |
| DDX58 | ENSG00000107201 | 9 | **4.06** | 3.43 | **3.32** | 2.97 |
| FKBP5 | ENSG00000096060 | 6 | **-5.44** | **-5.06** | **-4.93** | -3.59 |
| HMGB2 | ENSG00000164104 | 4 | **-4.01** | **-4.07** | **-3.56** | **-3.94** |
| KLF9 | ENSG00000119138 | 9 | **-4.86** | -3.62 | **-4.73** | -2.19 |
| LCP2 | ENSG00000043462 | 5 | **3.39** | 2.39 | **3.22** | 0.12 |
| NDUFAF5 | ENSG00000101247 | 20 | **-4.34** | -1.8 | **-3.65** | 0.48 |
| SESN3 | ENSG00000149212 | 11 | **-3.39** | **-3.89** | **4.63** | -1.96 |
| SH2D3C | ENSG00000095370 | 9 | **3.6** | 3.22 | **3.24** | 2.09 |
| SLA | ENSG00000155926 | 8 | **-4.01** | **-4.16** | **-3.99** | -2.79 |
| SNN | ENSG00000184602 | 16 | **-4.14** | -3.07 | **-3.47** | **-4.09** |
| TIPARP | ENSG00000163659 | 3 | **-3.68** | -3.54 | **-4.28** | -0.18 |
| TSC22D3 | ENSG00000157514 | X | **-3.98** | -3.15 | **-4.36** | -2.16 |
| TUBA4A | ENSG00000127824 | 2 | **-4.79** | -2.89 | **-3.97** | -2.14 |
| TXLNA | ENSG00000084652 | 1 | **4.16** | 3.06 | **4.19** | 2.37 |
| VSIR | ENSG00000107738 | 10 | **-4.42** | **-4.3** | **-4.27** | -3.67 |
| ZBTB16 | ENSG00000109906 | 11 | **-6.9** | **-4.45** | **-7.27** | **-4.12** |
| ZNF564 | ENSG00000249709 | 19 | **-3.98** | -3.17 | **-3.47** | -1.9 |

Bold font represents FDR-adjusted $p < 0.05$ .
PBMC samples are from the replication study.
HGNC: HUGO Gene Nomenclature Committee; Mono: monocytes; PBMC: peripheral blood mononuclear cells; Tmem: memory T cells; Tn: naive T cells.

Finally, we performed pathway enrichment analysis on the differentially expressed genes from each of the 6 cell-type-state contexts in order to identify shared and distinct pathways involved in immune dysregulation among T cells and monocytes in MS. The top enriched gene ontology (GO)^20^ biological processes (FDR-adjusted *p* <0.05) for T cells and monocytes were T cell activation and myeloid cell activation, respectively. No enriched GO pathways were found for genes from the stimulated monocytes. Additional pathway enrichment analysis using MsigDB hallmark gene sets^21^ identified significant enrichment for several hallmark pathways among the differentially expressed genes from Tn cells and monocytes (Figure 2F, Supplementary Table 3), several of which were previously implicated in MS^16,22^. However, our analysis provides further information on the cell-type-specificity of their involvement in MS pathology. Pathways such as PI3K/AKT/mTOR and mTORC1 signaling, inflammatory response, and transplant rejection were enriched among differentially expressed genes from both cell types. On the other hand, pathways such as IFN-γ response and the complement system were enriched among differentially expressed genes from Tn cells, but not monocytes; and genes from TNF-α / NF-κB signaling, IL-2 / STAT5 signaling, hypoxia, and glycolysis and gluconeogenesis pathways were enriched among genes from monocytes, but not Tn cells.

### T cell and monocyte co-expressed gene modules associated with MS

To complement our gene-level results, we elected to complete analyses after implementing a dimension reduction approach to help boost our statistical power to discover novel patterns of perturbed transcriptomic responses in MS. Specifically, we performed weighted gene coexpression network analysis (WGCNA)^23^ to (1) identify sets of co-expressed genes (gene modules) in each of the 6 cell-type-state transcription profiles, and (2) determine the association of the identified gene modules with MS. Given that biological processes often involve activation and/or suppression of several genes in concert with each other, the agnostic identification of groups of co-expressed genes in our data could potentially identify novel pathways that may be missing from the above-used predefined gene sets (i.e., GO biological processes and MsigDB hallmark).

Our WGCNA analysis was conducted separately for each cell-type-state context, and identified a total of 197 co-expressed gene modules from all 6 cell-type-states. Of note, our modules were set to include both positively-and negatively-correlated genes. The list of genes belonging to each module and the top 10 GO gene sets enriched in each module can be found in the Supplementary Data. We note that at least one identified module from each cell-type-state was highly associated with sex (Supplementary Table 4) and mainly consisted of known sex-associated genes from X and Y chromosomes. This observation supports the biological relevance of our identified modules.

Of our 197 gene modules, 6 modules were significantly associated with MS (Bonferroni-adjusted *p* <0.05, Figure 3A, Supplementary Table 4). The association analysis was performed between the eigengene from each module (i.e., the first principal component of the expression of the genes in each module) and MS diagnosis. The hub gene of each MS-associated module (i.e., a highly connected gene from each module, identified as part of the WGCNA analysis) is shown in Figure 3A and is used for module annotation throughout the rest of the manuscript. Secondary association analysis using module eigengenes demonstrated that the 6 MS-associated modules were significantly different in both untreated MS vs. healthy and GA-treated MS vs. healthy comparisons; and no module showed a significant difference between untreated and GA-treated MS participants (Supplementary Table 4). These findings are in line with our earlier observations from the transcriptome-wide association analysis that highlighted the similarity between the untreated and GA-treated MS transcriptomes.

**Figure 3.**
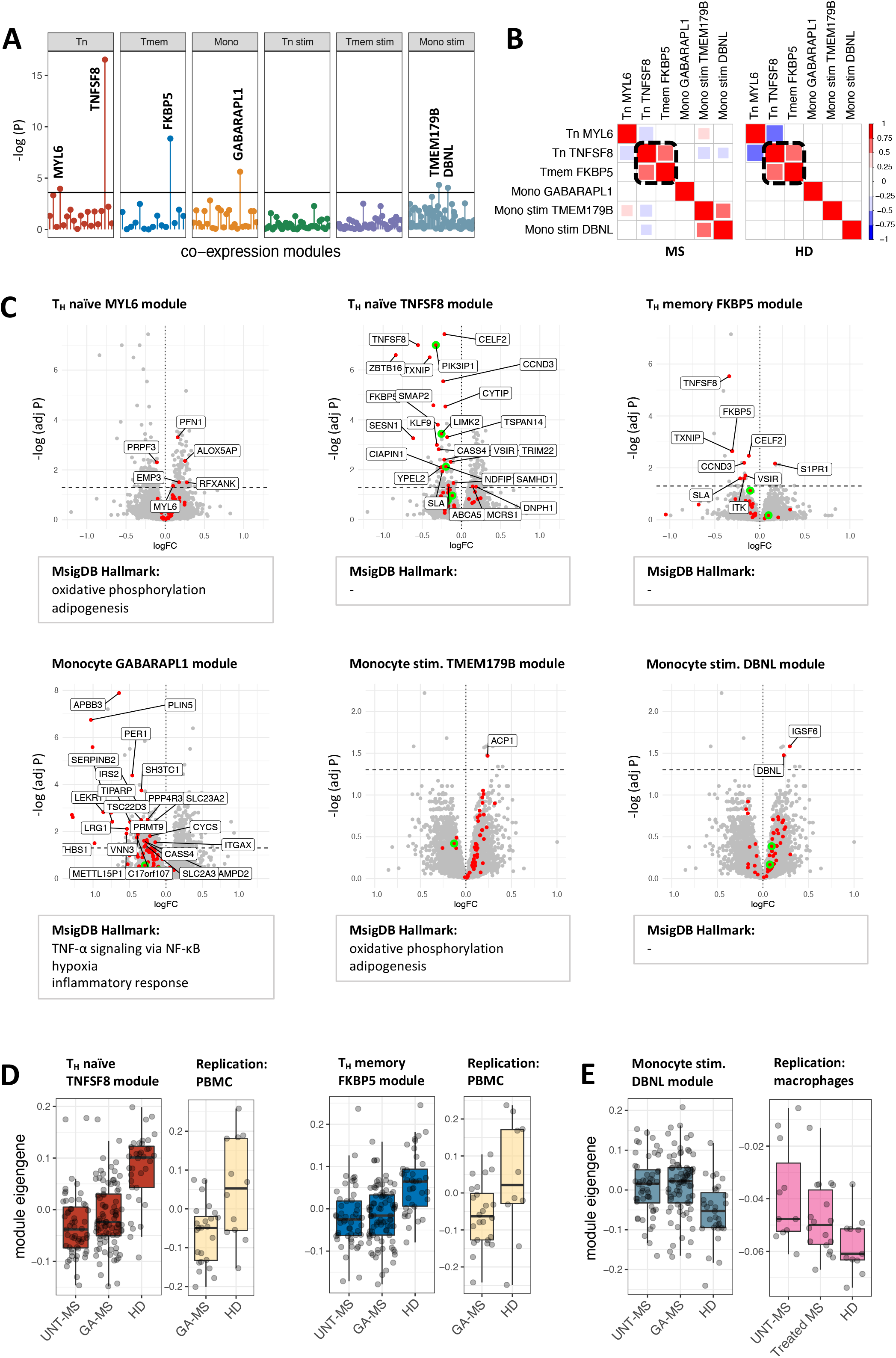
Results from weighted gene co-expression network analysis. (A) Co-expression modules associated with MS and their hub genes. Horizontal black line represents the significance level (Bonferroni-adjusted *p* =0.05). (B) Correlations between MS-associated module eigengenes in MS and healthy participants, separately. The two highly correlated modules in both MS and healthy individuals are highlighted using the dashed black squares. (C) Associations between the genes in each of the 6 MS-associated modules and MS are visualized using volcano plots. Red dots represent genes belonging to each module. Horizontal dashed line represents the significance level (FDR-adjusted *p* =0.05). Differentially expressed genes from each module are labeled in black. Green circles represent genes that are also associated with genetic susceptibility to MS (T_H_ naïve TNFSF8 module: *GGNBP2, LIMK2, NDFIP1, PIK3IP1, ZBTB38;* T_H_ memory FKBP5 module: *ETS1, NR1D1;* monocyte GABARAPL1 module: *CXCR4, MPV17L2;* stimulated monocyte TMEM179B module: *ECD;* stimulated monocyte DBNL module: *ANXA7, RNFT1).* Enriched MsigDB hallmark gene sets are also shown for each module. (D) Replication of the association between T cell TNFSF8/FKBP5 modules and MS using independent PBMC samples. (E) Replication of the association between stimulated monocyte DBNL module and MS using independent data from monocyte-derived macrophages.

Comparing the list of genes from the MS-associated modules, we found that the TNFSF8 module (n=54 genes) from unstimulated Tn cells and the FKBP5 module (n=30 genes) from unstimulated Tmem cells shared a significant number of genes (n=11). Correlation analysis between the module eigengenes further supported the similarity between these modules, as these modules were the only modules that were highly correlated (Spearman’s ρ >0.5, *p* <0.01) both among MS and healthy participants (Figure 3B). The two modules were the top associated modules with MS (*p* =2.8×10^-17^ and 1.4×10^-9^, respectively), and they were both enriched for genes from GO annotation’s T cell activation pathway.

Figure 3C further visualizes the association between individual genes from each MS-associated module and MS diagnosis (FDR-adjusted *p*-value and log fold change from the transcriptome-wide association analysis; differentially expressed genes are labeled). In addition to recapitulating some of the previous enrichment results from the differentially expressed genes, hallmark enrichment analysis on genes from the MS-associated modules provided support for the involvement of additional pathways such as oxidative phosphorylation and adipogenesis in MS. This suggests that although these pathways were not significantly enriched among the differentially expressed genes, the coordinated subthreshold change in the expression of their genes in MS could be detected using the co-expression analysis. Both of these pathways were previously implicated in MS pathophysiology^24–26^. Additional enrichment analysis using genes found in genetically-defined MS susceptibility loci^2^ revealed a nominal enrichment for MS susceptibility genes in the TNFSF8 module of Tn cells (*p* =0.02, Figure 3C), suggesting that the transcriptional changes in this module may provide a link between genetic susceptibility to MS and MS onset.

### Replication of module associations with MS in independent data

We used available RNA-seq data from independent PBMC and macrophage samples from MS and healthy participants in order to replicate the association between our modules and MS. Given that PBMC consist of both lymphocytes and monocytes, we used the PBMC data for replication analysis for all 6 MS-associated modules. However, the macrophage data were only used to assess replication for the 3 monocyte associations.

The PBMC data used for the replication study were part of the previously mentioned PBMC data used in the transcriptome-wide association analysis. However, we excluded participants that had samples in the 6 cell-type-state study so that only independent participants would be assessed in the replication analysis. Module eigengenes (i.e., the first principal component) were calculated and used for the association analysis. We found significant evidence of replication for the Tn TNFSF8 module (n=24 GA-treated MS vs. n=12 healthy participants, *p* =0.01). Additionally, we observed suggestive evidence that the Tmem FKBP5 module (n=24 GA-treated MS vs. n=10 healthy participants, *p* =0.09) may also replicate if larger sample sizes are assessed (Figure 3D).

The macrophage data used for the replication of the MS-associated monocyte modules were repurposed from a separate study of monocyte-derived macrophages, which was performed and analyzed by an independent set of investigators at the Paris Brain Institute^25^. In that published study, CD14^+^ monocytes from MS and healthy participants were isolated from peripheral blood and differentiated into macrophages through *in vitro* exposure to GM-CSF (details in the Methods, Table 1). The eigengene for the DBNL module found in stimulated monocytes in our data (Figure 3A) was also significantly differentially expressed in this macrophage dataset which consists of healthy (n=11) and both untreated (n=11) and treated (n=17) MS participants: *p* =0.02 for the comparison to untreated MS and *p* =0.05 for treated MS (Figure 3E).

Given our moderate sample sizes in these replication efforts and the fact that the cell types from both of our replication datasets were different from the cell types used in the discovery analysis, the other non-significant results do not rule out that the nonreplicated modules are truly perturbed in MS. Future dedicated larger studies will have to answer that question. However, our replication analyses did return significant results for certain modules in both T cells and monocytes; the T cell TNFSF8/FKBP5 modules and the monocyte DBNL module are therefore prioritized as signatures of transcriptional perturbation in the asymptomatic phase of MS (i.e., in clinical remission). They therefore offer tools for further investigation of the effects of immunomodulatory treatments now routinely used in MS on transcriptional signatures of MS-related immune perturbations.

### *In silico* drug screen for reversing MS-associated modules

We then queried the Connectivity Map (CMap)^27^ database to identify compound classes with reverse transcriptional signatures to each of the 6 MS-associated modules. CMap contains data from the transcriptional effects of ~5,000 small-molecule compounds, each tested in multiple cell lines (although none of which are of leukocyte origin). The top significant compound classes identified for each module are shown in Figure 4A. Intriguingly, glucocorticoid receptor agonists were found as the top compound class for the Tn TNFSF8 and Tmem FKBP5 modules. Among the prioritized compound classes, glucocorticoid receptor agonists are the only drugs that are currently used in clinical management of MS. Our finding suggests that this class of drugs might exert part of their therapeutic effect by upregulating these T cell modules that we identified and validated as being downregulated in MS.

**Figure 4.**
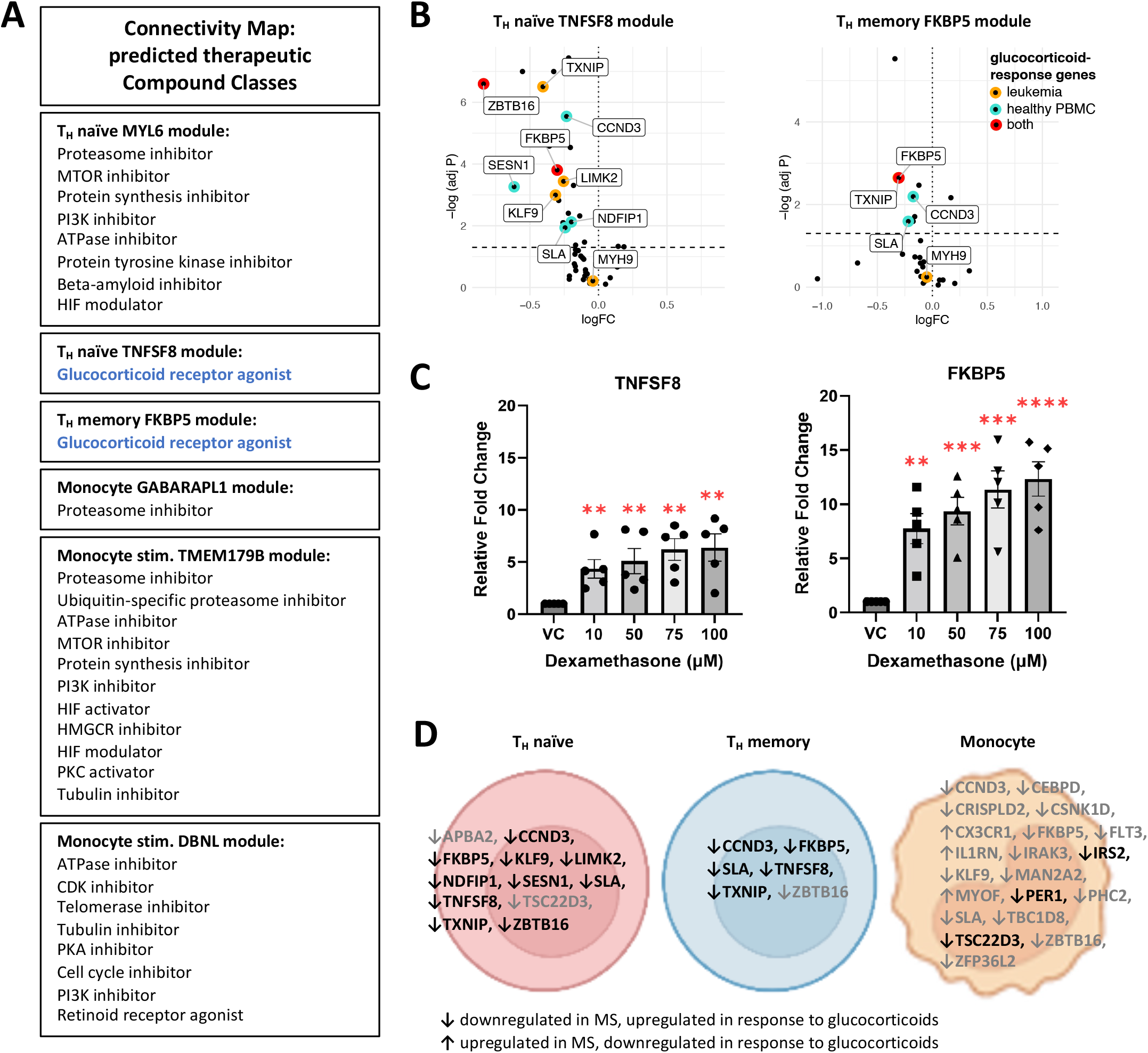
Relation between MS-associated genes/modules and glucocorticoid response genes. (A) Results from querying Connectivity Map database: top compound classes predicted to reverse changes in each MS-associated module. Glucocorticoid receptor agonists (highlighted in blue) are the only therapeutic class currently in clinical use. (B) Enrichment of the MS-associated T cell TNFSF8/FKBP5 modules for glucocorticoid response gene sets. Volcano plots show association between the genes in each module and MS diagnosis. Horizontal dashed line represents the significance level (FDR-adjusted *p* =0.05). Orange and blue circles represent glucocorticoid response genes in leukemic cells and healthy PBMC, respectively. Red circles represent genes shared between the two glucocorticoid response gene sets. (C) Increase in the expression of *TNFSF8* and *FKBP5* in MS PBMC in response to *in vitro* treatment with dexamethasone. ** *p* <0.01, *** *p* <0.001, **** *p* <0.0001. VC= vehicle control (DMSO, 0 μM dexamethasone). (D) Genes with opposite direction of effect in association with MS (FDR-adjusted *p* <0.05) and in response to glucocorticoids. Black font represents genes belonging to the identified MS-associated modules (T_H_ naïve TNFSF8, T_H_ memory FKBP5, and monocyte GABARAPL1 modules, respectively).

Given this finding, we searched the literature for studies on transcriptional effects of glucocorticoids. We took two gene sets from two separate studies^28,29^ on *in vitro* effects of the glucocorticoid drug prednisolone on gene expression in immune cells in order to assess the enrichment of our modules for glucocorticoid response genes. The two studies were performed using different cell types (primary leukemic cells from acute lymphoblastic leukemia participants^28^ and PBMC from healthy individuals^29^), and the two sets of differentially expressed genes after glucocorticoid treatment (n=51 and 82 genes, respectively) had only 6 genes in common (Supplementary Table 5). However, the enrichment analysis using both gene sets demonstrated similar results: among all 197 modules, Tn TNFSF8 and Tmem FKBP5 modules were the only modules with significant enrichment for glucocorticoid response genes (Bonferroni-adjusted *p* <0.05, Figure 4B). Combining the genes from the two sets resulted in even more significant results (*p* =2.5×10^-13^ and 5.9×10^-7^ for Tn TNFSF8 and Tmem FKBP5 modules, respectively). Among the MS-associated modules, we also found a nominal significance for enrichment in glucocorticoid response genes in monocyte GABARAPL1 module (*p* =0.004 for the combined gene set).

Given that MS relapses are treated with a pulse of glucocorticoids, the majority of MS participants have a positive history of previous glucocorticoid treatment. However, none of our MS participants had experienced a relapse in <3 months prior to blood sampling. Given that the biological half-life of glucocorticoids is <72 hours^30^, and the transcriptional effects of oral glucocorticoids are shown to reverse to baseline in <48 hours^29,31^, the perturbation of glucocorticoid response genes in our MS samples is unlikely to be a long-lasting effect of previous glucocorticoid treatment. This is further supported by a lack of significant difference in the Tn TNFSF8 and Tmem FKBP5 module eigengenes between MS participants who had a relapse in the last 3-12 months (n=16) vs. in >12 months (n=127) prior to blood sampling (*p* =0.68 and 0.84, respectively; Supplementary Figure 3). On the other hand, the fact that the genes in common between the MS-associated modules and the glucocorticoid response gene sets showed an opposite direction of effect (i.e., they were all downregulated in MS and upregulated in response to glucocorticoid treatment), suggests an intrinsic defect in glucocorticoid response in MS. This is in line with previous literature on the decreased glucocorticoid sensitivity observed in MS PBMC^32,33^.

### Validation of glucocorticoid effect on gene expression in MS

In the previous sections, our analyses identified genes belonging to MS-associated modules that are downregulated in MS and are shown to be upregulated in response to *in vitro* glucocorticoid treatment in two non-MS studies^28,29^. Given the differences between glucocorticoid response in leukemic cells^28^ and cells from healthy participants^29^, one can assume that glucocorticoid response in MS may also be to some extent different. Hence, we decided to assess the effect of glucocorticoid treatment on the expression of genes from Tn TNFSF8 and Tmem FKBP5 modules in samples from MS participants. Our experiment on the effect of *in vitro* exposure to the glucocorticoid drug dexamethasone on PBMC samples from MS participants (n=5) demonstrated a significant increase in the expression of T cell hub genes *TNFSF8* and *FKBP5* across all tested doses (all *p* <0.01, Figure 4C). Unlike *FKBP5, TNFSF8* was not part of our two glucocorticoid response gene sets^28,29^. However, the upregulation of both hub genes in response to dexamethasone in samples from MS participants provides support for the mechanism through which glucocorticoids might exert part of their therapeutic effect in MS: they may normalize the expression of these T cells modules that are under-expressed in the context of MS and thus reduce the likelihood of T cell activation.

### T cell and monocyte MS-associated glucocorticoid response genes

As a follow up analysis to our finding on the enrichment of glucocorticoid response genes in MS-associated modules, we sought to investigate whether similar enrichment could be observed in all MS-associated genes in T cells and monocytes. Our analysis revealed significant enrichment for glucocorticoid response genes (combined set, in addition to *TNFSF8*) among differentially expressed genes in all 3 primary cell types (*p* =9.9×10^-8^, 3.9×10^-6^, and 3.1×10^-11^ for Tn, Tmem, and monocytes, respectively). However, we observed an interesting difference in the pattern of enrichment between T cells and monocytes: the majority of the differentially expressed glucocorticoid response genes in T cells were co-expressed and belonged to a single MS-associated module, while this was not the case for monocytes (Figure 4D). This explains the lack of a prominent glucocorticoid responsive MS-associated module in monocytes, and on the other hand suggests that glucocorticoid treatment has a strong effect on a single co-regulated pathway in MS T cells. This observation further provides human support for results from a previous murine study^34^, in which the therapeutic effect of glucocorticoids was demonstrated to be dependent on the presence of glucocorticoid receptors in T cells but not myeloid cells.

### Gene and module associations with MS susceptibility, severity, and progression

In addition to investigating associations between individual genes and gene modules with MS diagnosis, we performed exploratory analyses within the MS participants in association with several MS-related variables: (1) MS susceptibility assessed using MHC, non-MHC, and total MS polygenic scores; (2) MS severity measured using the Expanded Disability Status Scale (EDSS), brain parenchymal fraction (BPF) from magnetic resonance imaging, and T2 hyperintense white matter lesion burden; and (3) MS progression assessed using change in EDSS over 4 years of follow up, and the slope of change in brain parenchymal fraction and white matter lesion volume. None of our 197 modules showed significant association with any of the outcomes (all Bonferroni-adjusted *p* >0.05). This might be explained by our small sample size for identifying factors driving the heterogeneity of disease course and the low variability within our MS participant population who were mostly in the early phase of their disease. However, our gene-level analyses identified a few significant associations (Supplementary Table 6). The most prominent associations were observed between the MS MHC polygenic score and the expression of MHC class II genes *HLA-DRB1* and *HLA-DRB5.* These associations were found in all cell types except for Tn cells, and are in line with previous literature on the effect of the MS top HLA risk allele HLA-DRB1*1501 on the expression of these genes^35^. Other associations were less significant and need to be replicated in independent MS datasets.

## Discussion

Our study of transcriptional profiles from three cell types under both unstimulated and stimulated conditions provides new insights into perturbation of transcriptional programs in certain immune cells implicated in MS. Using a relatively large cell-type-state-resolved transcriptomic dataset of MS and healthy participants (1,075 transcriptomic profiles from 209 individuals), we identified shared and distinct changes in individual genes, biological pathways, and co-expressed gene sets in MS T cells and monocytes. We replicated part of our results in independent data from PBMC and monocyte-derived macrophages. Further, we complemented our results by performing an *in silico* drug screen and validating its findings using independent enrichment analyses and *in vitro* experiments.

Our analyses demonstrated that primary T cells and monocytes purified from peripheral blood show largely distinguishing transcriptomic signatures in MS, supporting their use as accessible cell models for studying MS immunopathophysiology. On the other hand, despite the large number of MS-associated genes in primary, *ex vivo* T cells and monocytes, we did not find similarly large differences in our *in vitro* stimulated cell types in MS: few genes were found to be differentially expressed in MS in TNF-α stimulated monocytes, and none were in anti-CD3/CD28 stimulated T cells. This observation suggests that *in vitro* stimulation of T cells with anti-CD3 and anti-CD28 antibodies does not uncover MS-related transcriptional alterations: there does not appear to be a change in the functional capacity of these cells that can be elicited by these stimulation paradigms or measured by transcriptional profiling. Our data suggests that stimulation appears to suppress disease-associated signatures found *ex vivo*. We are limited to RNA sequence data, so it is possible that other functional assessments may be more informative. However, our results are informative for future study designs, suggesting that deeper *ex vivo* profiling may be more productive than investing effort and resources in developing and deploying more stimulation paradigms.

Our transcriptome-wide association studies on data from unstimulated cell types identified >200 MS-associated genes in T cells and >400 genes in monocytes. The majority of these genes show a cell-type-specific association with MS. However, results from pathway enrichment analysis suggest that in addition to affecting distinct pathways in different cell types, part of the MS-related T cell-and monocyte-specific perturbations converge on the same biological pathways across both cell types. These shared pathways include PI3K/AKT/mTOR and mTORC1 signaling^22^. A similar finding was observed between the MS-associated unstimulated T cell MYL6 module and stimulated monocyte TMEM179B module: despite sharing few genes, both modules are enriched for oxidative phosphorylation and adipogenesis pathways^24–26^. These observations demonstrate that shared pathways can be affected through distinct patterns of transcriptional changes across different cell types. One gene stands out in this regard, *ZBTB16*: it is downregulated in both T and myeloid cells, and the effect was validated in PBMC as well. These results in two different cell types prioritized this transcriptional repressor for us, and it joins a substantial literature that has implicated ZBTB16 in autoimmunity in the past: it was proposed as a hub gene in a type 1 diabetes PBMC-based network^19^ and was reported to be altered in expression in SLE as well^18^. Its binding site is disrupted by a thyroiditis susceptibility variant^36^, and while it has been studied more extensively in NK cells, it has been implicated in polarization of Th17 cells^37^, a cell type that plays an important role in MS autoimmunity.

Interestingly, it also appears to be involved in adipogenesis^38^ suggesting that it may be a link to obesity, which is a risk factor for MS. Overall, this literature and our results suggest that ZBTB16 may be an important regulator of autoimmunity and should be prioritized in future studies.

Our dimension reduction approach of identifying co-expressed gene modules complemented the results of our gene-wise studies and identified 6 MS-associated modules. Of these, we were able to offer evidence of replication in independent samples processed in a different manner for two T cell and one monocyte module, increasing our confidence of their relevance to MS. These statistically robust results – for the TNFSF8, FKBP5 and DBNL modules (Figure 3) – open several interesting avenues of investigation: they can be considered as biomarkers of MS. Our initial analyses of selected MS outcome measures suggests that these modules are not related to disease course, although larger datasets should be evaluated to produce more definitive results on that front. However, these biomarkers could be very informative in the diagnostic work-up for MS, providing some quantitative support to our traditional diagnosis of exclusion, and it would be interesting to see whether it correlates with other biomarkers of MS, such as the presence of plasma cells in the CSF^39,40^. At this time, we still do not have a reliable blood-based biomarker for MS. More importantly, the eigengenes of these 3 modules (the summary measures of each of the gene signatures) could prove even more useful in longitudinally monitoring individuals at risk for MS, as there is increasing interest in understanding the immune state of such individuals and to involve them in clinical trials. It is likely that a subset of at risk individuals are in an intermediate state, with evidence of peripheral immune dysfunction that has not yet engaged the target organ (in this case the brain) to cause disease. The fact that these signatures are present in PBMC also suggests that we may not need to purify cell types to assess the utility of these biomarkers: now that we have identified signatures, we can detect them in more complex mixtures such as PBMC or whole blood.

One limitation of our unbiased transcriptome-wide association-based study is that, while we now have robust observations, we do not yet know how these peripheral immune perturbations relate to the disease. Our analyses of glucocorticoid-related signaling illustrates one approach: to identify and test compounds that could correct the immune perturbation in humans *in vivo* and see whether this affects either (1) the onset of MS in someone at risk, (2) reduces the relapse rate, or (3) influences another outcome measure. The prioritization of glucocorticoids in our *in silico* analyses is an interesting proof of concept given their clinical utility in managing acute relapses; at the high doses typically used in pulse methylprednisolone treatments, it is likely that multiple molecular effects are occurring. We have now described one of them that seems to be clearly related to an altered immune response; however, the adverse effects of chronic steroid use suggest that it may be best to pursue some of the other therapeutic options that emerged from our *in silico* screen.

Impairment in the hypothalamic-pituitary-adrenal (HPA) axis has long been documented in MS: Individuals with MS have elevated serum adrenocorticotropic hormone (ACTH), increased cerebrospinal fluid and serum cortisol (the main human glucocorticoid), and impaired response to dexamethasone suppression and dexamethasone-suppressed corticotrophin-releasing hormone tests, coupled with decreased glucocorticoid receptor affinity and glucocorticoid sensitivity in PBMC^32,33,41^. These differences are found in all clinical subsets of MS, including individuals with primary-progressive disease course, who have no relapse and glucocorticoid treatment history^32^. The downregulation of glucocorticoid response genes in MS T cells and monocytes observed in our study may be explained by the decreased sensitivity to glucocorticoids. However, this needs to be validated in future studies where HPA axis hormones and glucocorticoid sensitivity are measured simultaneous with immune cell transcriptomes. The exact mechanisms underlying the decreased glucocorticoid sensitivity are also unclear. Previous studies have suggested that a decrease in the number of glucocorticoid receptors is unlikely to be the cause, as the density of glucocorticoid receptors is not affected in MS PBMC^32^. Similarly, the expression of *NR3C1*, the gene encoding the glucocorticoid receptor, was not different in our MS samples. However, we observed differential expression in a number of other genes involved in the formation or function of the glucocorticoid receptor complex: *HSP90B1* (encoding a chaperone for glucocorticoid receptor) in upregulated in MS T cells, *FKBP5* (encoding a co-chaperone of HSP90) in downregulated in MS T cells and monocytes, and *HMGB2* (encoding a non-histone chromosomal protein which facilitates DNA binding of glucocorticoid receptor) is downregulated in MS T cells, monocytes, and PBMC. Whether these changes are linked to the mechanism underlying decreased glucocorticoid sensitivity in MS needs to be investigated in future studies.

Although our work benefited from a relatively large sample size, minimization of treatment confounding effects, cell type- and state-specific data generation, complementary analytical approaches, and replication and validation experiments, our study had several limitations, some of which were discussed above. These included the limitations of the studied *in vitro* stimulation models and the unavailability of measures assessing HPA axis impairment and MS activity. Another major limitation of our study was that it did not include data on other peripheral immune cell types implicated in MS pathology such as B cells, plasma cells, and cytotoxic T cells^16^. Further, the use of bulk RNA-seq data from purified helper T cells and classical monocytes might have masked more fine-grained associations that could potentially be observed in subtypes of these cells. Technological advances helping with feasibility of large-scale and high-resolution single-cell RNA-seq studies can alleviate both of these issues in future investigations. Finally, the enrichment of the glucocorticoid-responsive T cell TNFSF8 module for MS susceptibility genes suggests a causal role for glucocorticoid response dysfunction in MS pathology. However, additional studies are required to elucidate the cascade of pathological vs. compensatory molecular events involved in glucocorticoid response pathology in MS, and to elaborate on the serial vs. parallel nature of transcriptional changes observed in different cell types.

Notwithstanding the limitations, our study has helped resolve cell-type-specific perturbations in transcriptomic programs in MS helper T cells and monocytes. Additionally, our approach of combining data-driven identification of coordinated transcriptional changes with *in silico* drug screen has helped provide a novel explanation for the therapeutic effect of glucocorticoids in connection with MS T cell pathology. Our dataset can continue to be used by the scientific community as a resource for replication or meta-analysis studies, or further analyzed in conjunction with other molecular datasets for hypothesis testing or discovery.

## Methods

### Participants

Multiple sclerosis (MS) participants were from the Comprehensive Longitudinal Investigation of Multiple Sclerosis at the Brigham and Women’s Hospital (CLIMB) study^11^. CLIMB is a natural history observational study of MS, in which participants undergo semi-annual neurological examinations and annual magnetic resonance imaging (MRI). The participants also donate blood, from which peripheral blood mononuclear cells (PBMC) are extracted and cryopreserved. In 2016-2017, cryopreserved PBMC samples from participants meeting the following criteria were pulled from the archive for sample processing: (1) age 18-60 years old at the time of sampling, (2) a diagnosis of MS based on the 2010 McDonald criteria, (3) a relapsing-remitting disease course, (4) being untreated or on treatment with glatiramer acetate (GA) for >6 months at the time of sampling, (5) no evidence of disease activity in the 6 months prior to sampling, (6) no steroid use in the 30 days preceding sampling, and (7) an Expanded Disability Status Scale (EDSS) score between 0 to 3 at the time of sampling.

Healthy participants were from the PhenoGenetic Project, a resource previously used to study the extent of variation in immune function among healthy individuals in the ImmVar study^13,14^. PBMC samples were from participants 18-60 years old who self-reported to be free of chronic infectious, inflammatory, and metabolic disease.

Both studies were approved by the Institutional Review Board of the Brigham and Women’s Hospital, and all participants had signed a written informed consent. PBMC samples from 227 participants who fulfilled the criteria (181 MS and 46 healthy participants) were used for further processing. 90% of samples were collected between 2008-2014.

### PBMC sampling

PBMC were collected prospectively from participants using the Immune Tolerance Network (https://www.immunetolerance.org/) protocol to ensure data quality and to minimize variation among samples. In short, fresh blood was processed within 4 hours using the Ficoll procedure to extract PBMC. The PBMC were then resuspended in fetal bovine serum with 10% DMSO and cryopreserved in liquid nitrogen.

### T cell and monocyte isolation and stimulation

PBMC samples meeting study criteria were pulled from the sample archive and thawed in batches by brief immersion in +37C water bath. Cell viability was assessed by acridine orange/propidium iodine staining and samples with <80% viability were discarded. Monocytes were pre-purified using CD14^+^ magnetic beads (Miltenyi) after which viable CD14^+^CD16^-^ monocytes were isolated from the positive fraction on a BD FACSAria flow cytometer using fluorochrome labelled antibodies: anti-CD3 FITC (clone: HIT3a; 1:50 dilution, 2 ml to 100 ml staining volume containing 1 × 10^6^ cells), anti-CD14 PE (M5E2; 1:50 dilution), anti-CD16 APC (3G8; 1:100 dilution), and 7-AAD (1:50 dilution). Viable naïve and memory CD4^+^ T cells were isolated from the negative fraction using anti-CD3 FITC, anti-CD4 PE-Cy7 (RPA-T4; 1:50 dilution), anti-CD45RA AF700 (HI100; 1:33 dilution), and 7-AAD. Cells for unstimulated conditions were sorted directly into lysing buffer (Norgen Biotek Corp) containing 2-Mercaptoethanol and frozen. Cells for stimulated conditions were cultured in serum-free X-Vivo medium using 2 mg/mL plate bound anti-CD3 (clone OKT3) and 1 mg/mL anti-CD28 (clone CD28.2) antibodies in the presence of 0.5 mg/mL rhIL-2 (Ro 23-6019, National Cancer Institute Biologics Repository) for T cells and using 10 ng/mL rhTNF-α (210-TA, R&D Systems) for monocytes. Stimulated monocytes were lysed and frozen after 4 hours. Stimulated T cells were stained with 7-AAD after 16 hours and viable cells were sorted directly into lysis buffer using a BD FACSAria flow cytometer.

### RNA-seq data generation and preprocessing

RNA was extracted from all primary and *in vitro* stimulated samples using an RNA/DNA purification kit (Norgen Biotek Corp) and processed using the Smart-Seq2 protocol^42^ to produce cDNA. Whole transcriptome 25-bp paired-end sequencing was performed for 1,176 samples from 227 individuals using the Broad Institute HiSeq 2500 platform to an average depth of 15 million reads per sample. Processing was performed according to the Broad Institute RNA-seq pipeline for the GTEx Consortium^43^. Briefly, RNA sequence reads were aligned to the GRCh38/hg38 genome reference using STAR^44^, quality control was performed using RNA-SeQC^45^ and quantification of gene expression levels were performed using RSEM^46^.

Prior to grouping samples from each cell-type-state for the rest of the preprocessing and analyses, data from participants who were from non-European ancestry (determined based on self-reported ethnicity and/or genetic data; details below; n=17) or had incomplete or mismatching demographic information (n=1) were excluded. This resulted in the retention of 1,075 transcriptome data from 209 individuals for further processing. Subsequent processing was performed on data from each group, separately, as follows. Transcripts with low expression values (average TPM <2) were removed, with expression data from an average of 11,714 genes per cell-type-state remaining for analysis. TPM values were log-transformed and quantile-normalized. Samples with outlier principal component (PC) values (>3 standard deviation on PCs that explained >5% of the variance) were further excluded. Expression levels were adjusted for the effects of technical confounding variables, i.e., library preparation batch and RNA-seq quality metrics that were associated with the first 10 PCs of expression data, such as the number of aligned reads, percentage of coding, intronic, and ribosomal bases, median coverage variability, and median 3’ and 5’ biases. Calculated residual expression values were used for the association analyses.

### Genetic data preprocessing and imputation

Genome-wide genotyping data were available for 218 out of 227 participants as part of larger batches of previously genotyped CLIMB and ImmVar samples. Genotyping was performed using Illumina MEGA^EX^, Infinium OmniExpressExome, and Affymetrix 6.0 arrays. Preprocessing of the genetic data was performed using PLINK^47^. SNPs with call rate <95% were removed. Multidimensional scaling was performed using HapMap3 reference data. Samples from individuals of non-European ancestry and ethnic outliers with >3 standard deviation difference from the European samples were identified and both their RNA-seq and genetic data were removed (n=17). Additionally, genetic data from samples with genotyping rate <90%, mismatch between recorded sex and genetic sex, and low or high heterozygosity rates (>3 standard deviation) were excluded (n=4). No remaining samples were from related individuals (all pi-hat <0.125). Imputation was performed for each genotyping batch separately. Wholegenome imputation was performed using the Michigan Imputation Server^48^ and the Haplotype Reference Consortium (HRC) panel v1.1. Imputed data from the study participants were then merged, and SNPs with low imputation quality (R^2^ <0.8) were removed. SNPs, amino acids and HLA alleles in the MHC region were imputed using SNP2HLA^49^ and the Type 1 Diabetes Genetics Consortium (T1DGC) HLA reference panel.

### MS susceptibility polygenic scores

MS polygenic scores were calculated as described previously^50^. Briefly, genome-wide significant results from the 2019 International Multiple Sclerosis Genetics Consortium (IMSGC) MS genome-wide association study^2^ (or if not available, tag SNPs in LD >0.8) were used for polygenic score calculation. MS MHC polygenic scores were calculated as the sum of the imputed dosages of 31 MHC markers associated with MS multiplied by their effect sizes (i.e., log odds ratio). Similarly, MS non-MHC polygenic scores were weighted scores calculated from the imputed dosages of 193 autosomal non-MHC MS-associated SNPs (7 loci did not have high-quality tag SNPs in our imputed dataset, and therefore were not included in polygenic score calculation). Finally, MS total polygenic scores were calculated as the sum of MHC and non-MHC polygenic scores.

### EDSS outcomes

179 MS participants had available EDSS data within 3 months of the blood sampling. From these participants, 132 were followed up for ≥4 years after the blood sampling visit and had at least 4 available longitudinal EDSS measurements (median=21 data points per person including visits prior to the time of blood sampling). In order to overcome the EDSS measurement variability and minimize the effect of temporary outlier measurements, smoothing splines were fit to longitudinal EDSS data using R smooth.spline function, with equivalent degree of freedom set to 4 (Supplementary Figure 4). EDSS progression was defined as the difference between smoothed EDSS measurements at year 4 and the time of blood sampling.

### MRI outcomes

Proton density-, T2-and T1-weighted brain MRI was performed for MS participants. MS lesions and brain tissue compartments (gray matter, white matter, and cerebrospinal fluid) were segmented using template-driven segmentation and partial volume artifact correction (TDS+)^51^ method. Results underwent quality control and manual correction where necessary. Brain parenchymal fraction (BPF) was calculated as the sum of gray and white matter volumes divided by the intracranial volume (i.e., the sum of gray matter, white matter, and cerebrospinal fluid volumes).

120 MS participants had available MRI data acquired using 1.5T scanners within 1 year of the blood sampling (MRI performed using 3T scanners were few and were excluded from this study). BPF and log-transformed lesion volume from these participants were used as surrogate quantitative markers for brain atrophy and white matter demyelination, respectively. From these participants, 93 had at least 4 available longitudinal MRI data acquired using 1.5T scanners during their follow-up period (median=6 data points per person including visits prior to the time of blood sampling). The rate of progression in brain atrophy and white matter demyelination were calculated for these participants as the linear slope of change in BPF and log-transformed lesion volume over the follow-up period, respectively (Supplementary Figure 4).

### Transcriptome-wide association studies

Transcriptome-wide differential expression analyses between diagnostic and treatment groups were performed using calculated residual expression values (details above) and limma^52^ robust regression modeling (robust lmFit and robust eBayes functions). All comparisons included age and sex as covariates. Adjustment for multiple comparisons was performed for the genes tested in each cell-type-state. Associations with false discovery rate (FDR)-adjusted *p* <0.05 in each comparison were considered significant.

Within-MS transcriptome-wide association studies were performed using the following quantitative outcome variables: MHC, non-MHC, and total MS susceptibility polygenic scores; baseline EDSS, BPF, and lesion volume; and longitudinal change in EDSS, and the rate of change in BPF and lesion volume. Outlier outcome variables were winsorized in case of |>3| standard deviation difference from the mean. Association analyses included additional covariates depending on the outcome variable: all within-MS associations accounted for the effects of disease duration and treatment group, in addition to age and sex; association analyses on longitudinal outcomes (i.e., 4-year change in EDSS, and the rate of change in BPF and lesion volume) additionally accounted for the effects of their related baseline measurements. FDR-adjusted *p*-values from each transcriptome-wide study underwent an additional level of adjustment for the 9 tested outcomes for each cell-type-state.

### Co-expressed gene modules identification and analyses

R package WGCNA^23^ (weighted gene co-expression network analysis) was used to identify modules of co-expressed genes using calculated residual expression values from each cell-type-state. WGCNA partitions the transcriptome into modules of highly co-expressed genes, where each gene is either part of a co-expression module or stays unassigned. Therefore, not all genes are assigned a module, and each gene can only belong to one module. Biweight midcorrelation (bicor) with outlier removal >5 percentile was used to avoid sensitivity to outlier expression values. Modules were set to be unsigned (i.e., contained both positively and negatively correlated genes). Minimum module size (number of genes in each identified module) was set to 20. Module eigengenes (first principal component of the genes belonging to each module) were calculated using WGCNA and used for association analyses with MS diagnosis and MS-related outcomes. Associations were assessed using robust regression modeling. Similar to the transcriptome-wide association studies, quantitative MS-related outcomes were winsorized when necessary, and covariates were included in the model depending on the outcome variable. Adjustment for multiple comparisons was performed using Bonferroni method for the tested modules for each outcome.

### Gene set enrichment analyses

Gene ontology (GO)^20^ and MsigDB hallmark^21^ gene set enrichment analyses were performed using FUMA^53^. Minimum overlap with each gene set was set to 3 genes, and results were reported at 5% FDR. Enrichment analyses for the other gene sets (i.e., glucocorticoid response and MS susceptibility-associated genes) were performed using hypergeometric test. The two sets of glucocorticoid response genes were obtained from ^28,29^. Genes associated with MS susceptibility loci were obtained from ^2^.

### Compound screen using Connectivity Map (CMap)

The Connectivity Map (CMap)^27^ database contains data on the transcriptional effects of ~5,000 small-molecule compounds, each tested in multiple cell lines. The database can be queried using the CLUE (CMap and LINCS Unified Environment, https://clue.io/) platform. We queried the database (v1.3) using genes from each MS-associated module to identify compound classes with reverse transcriptional similarity to each module. Results with FDR-adjusted *p* <1×10^-4^ are reported.

### Replication dataset #1: PBMC RNA-seq

Similar to the main study, PBMC samples used in this dataset (n=65) were from CLIMB (n=29) and PhenoGenetic (n=36) participants. However, they were retrieved and sequenced separately as part of a larger batch for a different PBMC study. The inclusion/exclusion criteria were similar to the main study, except that the MS participants were all GA-treated. RNA extracted from these samples underwent the same sequencing and preprocessing pipelines as the above. Age and sex were used as covariates for all association analyses. Transcriptome-wide differential expression analysis between MS and healthy participants was performed using the above-outlined limma pipeline. Eigengene association studies were performed on the subset of samples that did not have paired data with the main study. Linear regression modeling was used to compare eigengene values between diagnostic groups.

### Replication dataset #2: macrophage RNA-seq

Data used in this dataset (n=39) were from a study on monocyte-derived macrophages, which was performed and analyzed by independent investigators^25^. Methods for this study are described in details elsewhere^25^. Briefly, the study was approved by the French Ethics committee, and written informed consent was obtained from all study participants. Recruited MS participants fulfilled the diagnostic criteria for MS, and none of the participants had any other inflammatory or neurological disorders. CD14^+^ monocytes were isolated from the peripheral blood of MS and healthy participants and differentiated into macrophages through *in vitro* exposure to granulocyte macrophage colony-stimulating factor (GM-CSF, 500 U/ml, ImmunoTools). Cells were lysed after 96 hours, and extracted RNA underwent 75-bp single-end sequencing using a NextSeq 500 sequencer to an average depth of 60 million reads per sample. Data quality control was performed using fastp^54^, alignment was performed to the GRCh38/hg38 genome reference using STAR^44^, and gene expression quantification was performed using RSEM^46^. Sequencing batch effects were corrected using the removeBatchEffect function of the limma^52^ package, and corrected TPM values were used in the replication analyses. Wilcoxon non-parametric sign rank test was used to compare eigengene values between groups.

### Dexamethasone experiment

*In vitro* experiments were conducted on PBMC samples from MS participants recruited at the Columbia University MS Center (n=5; in remission). Blood sampling and PBMC isolation were performed using the same protocol as the main study. The isolated cells were incubated for 24 hours with dexamethasone (MicroSource Discovery System, Cat# 01500203) at five concentrations: 0, 10, 50, 75, or 100 μM in DMSO. RNA was isolated from the cells using RNeasy plus micro kit (QIAGEN, Cat# 74034) according to the manufacturer’s protocol, and reverse transcribed to cDNA using iScript cDNA synthesis kit (Bio-Rad, Cat# 1708891). TaqMan gene expression assays HS00174286_m1 and HS01561006_m1 were used to quantify the expression of *TNFSF8* and *FKBP5* genes in each sample. Expression levels were normalized to the expression of the housekeeping gene *B2M.* Quantitative RT-PCR was performed using an Applied Biosystems QuantStudio 3 Real-Time PCR System. Differences in gene expression between dexamethasone-treated samples and the vehicle control group were assessed using *t*-test.

### Statistical analysis

All statistical tests are described in their respective methods sections. All reported *p*-values are two-sided.

## Data availability

The RNA sequencing data (1,075 samples from 6 cell-type-states from 209 participants) will be deposited in the Synapse database under accession code xx [link]. As sensitive human data, these data can be accessed upon request, following the establishment of a Data Use Agreement with Brigham and Women’s Hospital, Dr. Howard Weiner. Supplementary Data will be deposited to the Synapse database under accession code xx [link] and will have public access.

## Acknowledgements

We are grateful to the participants for their invaluable contribution to the study. This work was supported by funding from Genzyme. Funding sources had no role in the analysis, interpretation of results or preparation of the manuscript.

DF is supported by the Koerner Family Foundation New Scientist Award, the Krembil Foundation, the Canadian Institutes of Health Research, the Canadian Foundation for Innovation, and the CAMH Discovery Fund. NAP was supported in part by National Multiple Sclerosis Society (grants JF-1808-32223 and RG-1707-28657).

## Author contributions

VKK, HLW and PLD contributed to the study design. PLD and PK supervised the data generation. CM and PN contributed to the data generation. HK, ISV, DH and NAP contributed to the RNA-seq data pre-processing and quality control. TR, HK and DF contributed to data analysis. TR, HK and PLD contributed to the analysis plan. AS performed the dexamethasone experiment and analyzed its data. HT contributed to the dexamethasone experiment. VZ supervised the generation and analysis of the macrophage RNA-seq data. JF contributed to the macrophage RNA-seq replication study. CLIMB study data acquisition was supervised by HLW. PhenoGenetic study data acquisition was supervised by PLD. TR and PLD contributed to the interpretation of results and drafted the manuscript. All authors read and revised the manuscript. PLD supervised the entire study.

## Competing interests

Authors declare no competing interests in relation to the study.

NAP is currently an employee of Novartis Institutes for BioMedical Research.

---

**Supplementary Figure 1.**
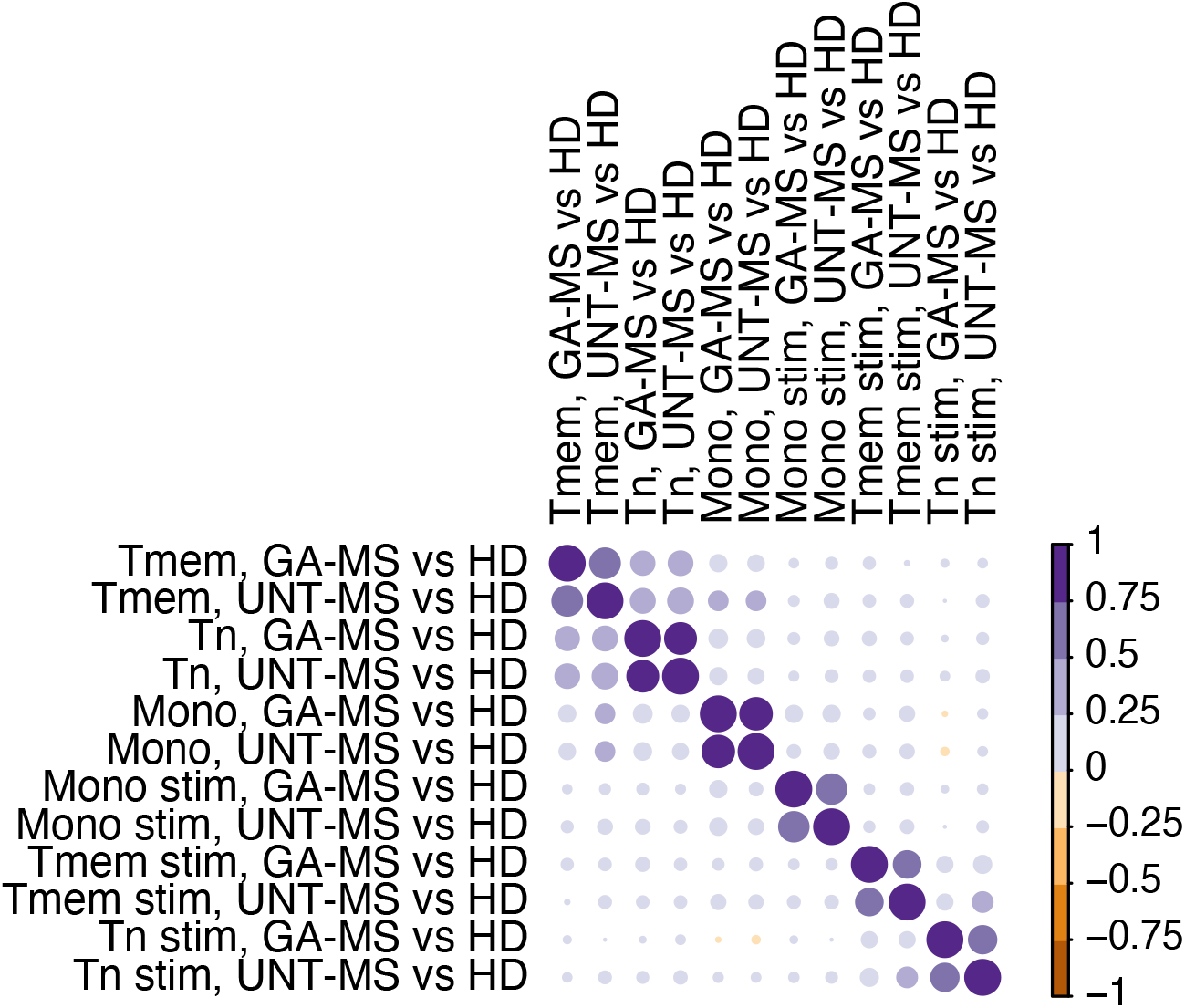
Correlations between *t*-statistics from the transcriptome-wide association studies comparing the 2 MS groups with healthy individuals.

**Supplementary Figure 2.**
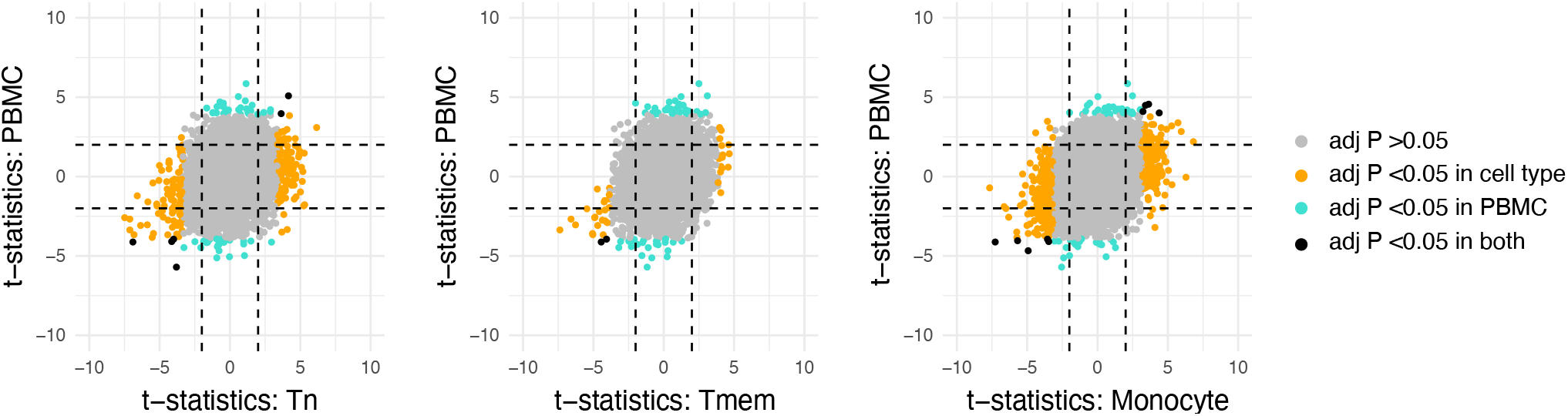
Correlations between *t*-statistics from the transcriptome-wide association studies comparing MS and healthy individuals in T cells and monocytes vs. PBMC. Orange dots: MS-associated genes in T cells or monocytes (FDR-adjusted *p* <0.05). Blue dots: MS-associated genes in PBMC (FDR-adjusted *p* <0.05). Black dots: MS-associated genes shared between the two comparisons. Horizontal and vertical dashed lines: nominal significance level (*p* =0.05). Spearman’s p from left to right: 0.17, 0.18, and 0.26, respectively.

**Supplementary Figure 3.**
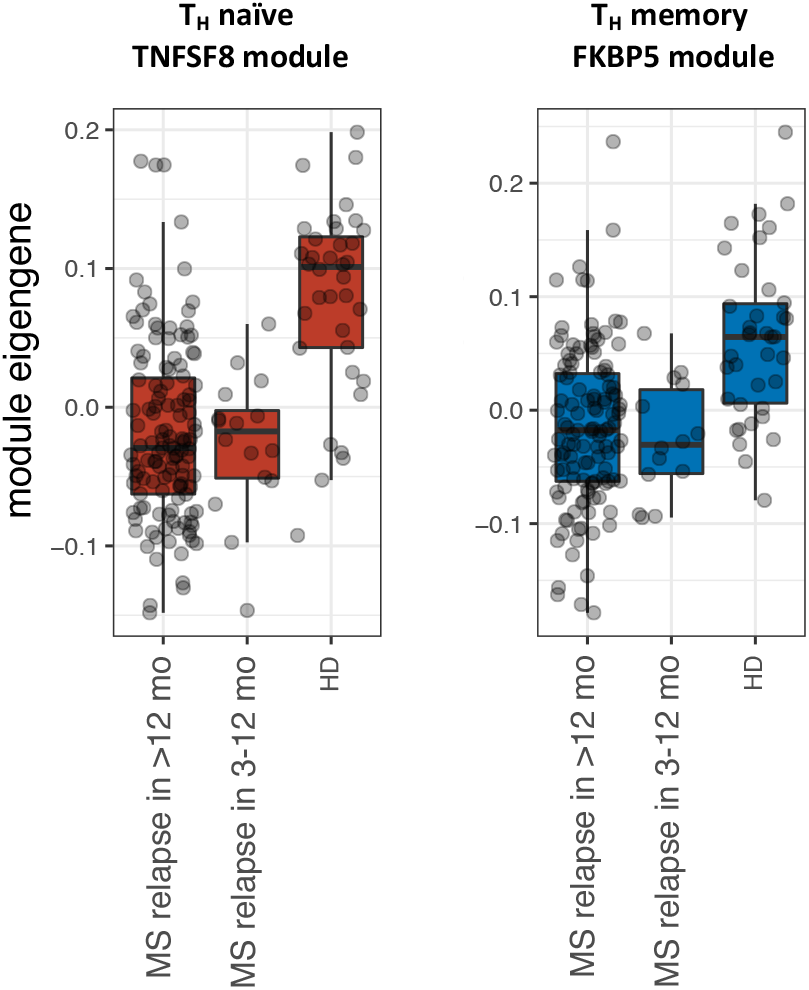
No difference in the T cell TNFSF8/FKBP5 module eigengenes between MS participants who had a relapse in the last 3-12 months vs. in >12 months prior to blood sampling.

**Supplementary Figure 4.**
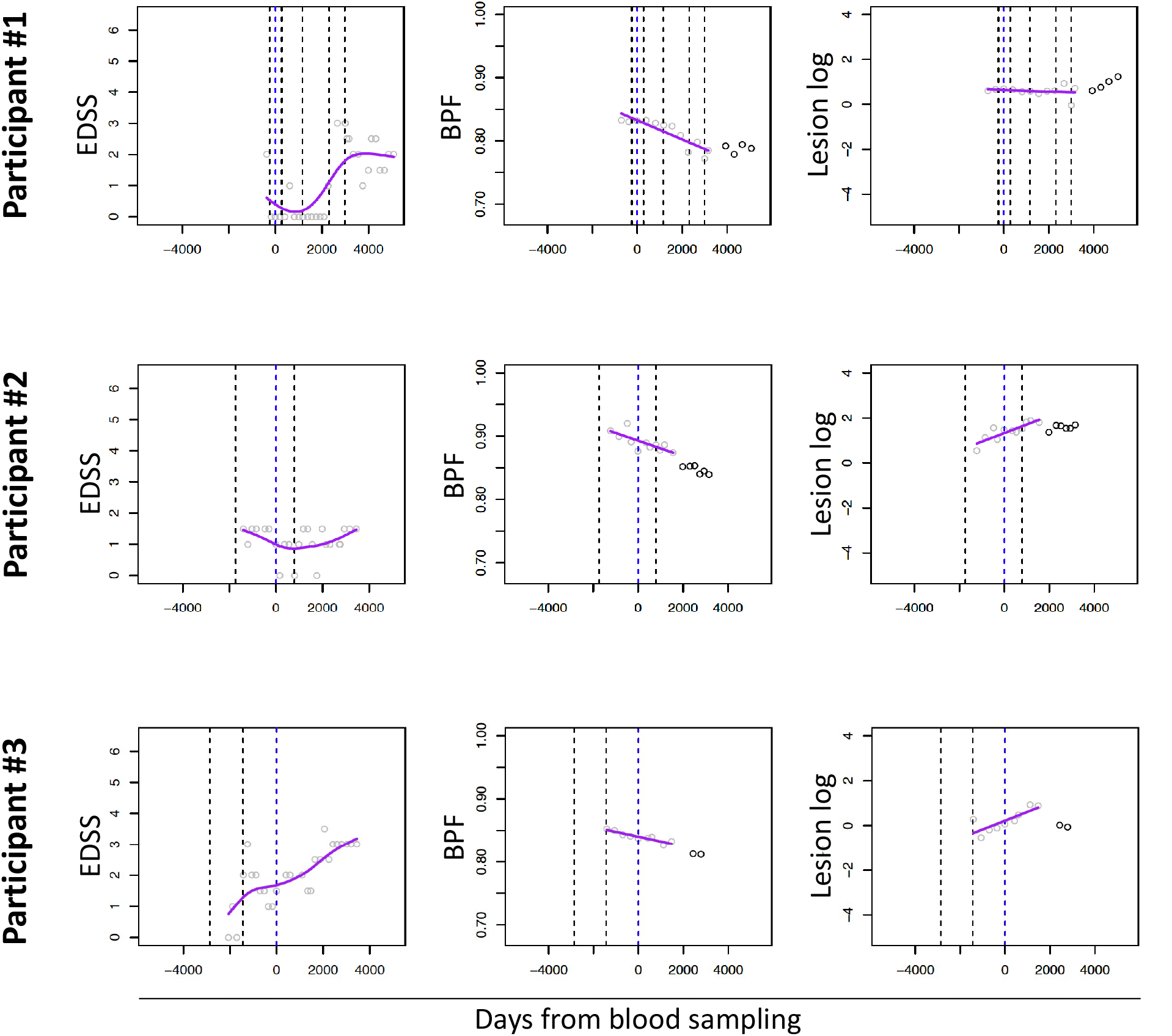
Example changes in EDSS, BPF, and log-transformed lesion volume over the follow up period are visualized for three of the MS participants. Circles represent individual measurements. MRI measurements shown in gray are from 1.5T MR scans. X-axis units are in days. Purple lines represents the fitted models. Vertical blue lines represent the time of blood sampling. Vertical black lines represent clinical relapses.

**Supplementary Table 1.** Genes differentially expressed between untreated and GA-treated MS.

**Supplementary Table 2.** Genes differentially expressed between MS and healthy participants in at least one of the six cell-type-states.

**Supplementary Table 3.** Enrichment of MS-associated genes in T cells and monocytes for MsigDB hallmark gene sets.

**Supplementary Table 4.** Associations between module eigengenes and outcome variables.

**Supplementary Table 5.** Glucocorticoid response genes.

**Supplementary Table 6.** Genes associated with MS outcome variables in each of the six cell-type-states.

## Notes

### Competing Interest Statement

The authors have declared no competing interest.

### Summary of Updates

Text and Figures are updated.

## References

1. Lassmann, H. Pathogenic mechanisms associated with different clinical courses of multiple sclerosis. Front. Immunol. 10, (2019).

2. International Multiple Sclerosis Genetics Consortium. Multiple sclerosis genomic map implicates peripheral immune cells and microglia in susceptibility. Science 365, eaav7188 (2019).

3. Reich, D. S., Lucchinetti, C. F. & Calabresi, P. A. Multiple Sclerosis. N. Engl. J. Med. 378, 169–180 (2018).

4. Nickles, D. et al. Blood RNA profiling in a large cohort of multiple sclerosis patients and healthy controls. Hum. Mol. Genet. 22, 4194–4205 (2013).

5. Gandhi, K. S. et al. The multiple sclerosis whole blood mRNA transcriptome and genetic associations indicate dysregulation of specific T cell pathways in pathogenesis. Hum. Mol. Genet. 19, 2134–2143 (2010).

6. Ratzer, R. et al. Gene expression analysis of relapsing-remitting, primary progressive and secondary progressive multiple sclerosis. Mult. Scler. 19, 1841–1848 (2013).

7. Brorson, I. S. et al. No differential gene expression for CD4 + T cells of MS patients and healthy controls. Mult. Scler. J. – Exp. Transl. Clin. 5, 205521731985690 (2019).

8. Kim, K. et al. Cell type-specific transcriptomics identifies neddylation as a novel therapeutic target in multiple sclerosis. Brain 144, 450–461 (2021).

9. Ingelfinger, F. et al. Twin study reveals non-heritable immune perturbations in multiple sclerosis. Nature 603, 152–158 (2022).

10. Gresle, M. M. et al. Multiple sclerosis risk variants regulate gene expression in innate and adaptive immune cells. Life Sci. alliance 3, (2020).

11. Gauthier, S. A., Glanz, B. I., Mandel, M. & Weiner, H. L. A model for the comprehensive investigation of a chronic autoimmune disease: the multiple sclerosis CLIMB study. Autoimmun. Rev. 5, 532–6 (2006).

12. Ottoboni, L. et al. An RNA Profile Identifies Two Subsets of Multiple Sclerosis Patients Differing in Disease Activity. Sci. Transl. Med. 4, 153ra131–153ra131 (2012).

13. De Jager, P. L. et al. ImmVar project: Insights and design considerations for future studies of “healthy” immune variation. Semin. Immunol. 27, 51–7 (2015).

14. Raj, T. et al. Polarization of the effects of autoimmune and neurodegenerative risk alleles in leukocytes. Science 344, 519–23 (2014).

15. Trickett, A. & Kwan, Y. L. T cell stimulation and expansion using anti-CD3/CD28 beads. J. Immunol. Methods 275, 251–255 (2003).

16. Dendrou, C. A., Fugger, L. & Friese, M. A. Immunopathology of multiple sclerosis. Nat. Rev. Immunol. 2015 159 15, 545–558 (2015).

17. Souren, N. Y. et al. DNA methylation signatures of monozygotic twins clinically discordant for multiple sclerosis. Nat. Commun. 10, (2019).

18. Fan, H. et al. Gender differences of B cell signature related to estrogen-induced IFI44L/BAFF in systemic lupus erythematosus. Immunol. Lett. 181, 71–78 (2017).

19. Jia, X. et al. Integrated analysis of different microarray studies to identify candidate genes in type 1 diabetes. J. Diabetes 9, 149–157 (2017).

20. Ashburner, M. et al. Gene ontology: tool for the unification of biology. The Gene Ontology Consortium. Nat. Genet. 25, 25–29 (2000).

21. Liberzon, A. et al. The Molecular Signatures Database (MSigDB) hallmark gene set collection. Cell Syst. 1, 417–425 (2015).

22. Wu, T. & Mohan, C. The AKT axis as a therapeutic target in autoimmune diseases. Endocr. Metab. Immune Disord. Drug Targets 9, 145–150 (2009).

23. Langfelder, P. & Horvath, S. WGCNA: an R package for weighted correlation network analysis. BMC Bioinformatics 9, 559 (2008).

24. La Rocca, C. et al. Immunometabolic profiling of T cells from patients with relapsing-remitting multiple sclerosis reveals an impairment in glycolysis and mitochondrial respiration. Metabolism. 77, 39–46 (2017).

25. Fransson, J. et al. Dysregulated functional and metabolic response in multiple sclerosis patient macrophages correlate with a more inflammatory state, reminiscent of trained immunity. bioRxiv 2021.01.13.426327 (2021) doi:10.1101/2021.01.13.426327.

26. Matarese, G., Carrieri, P. B., Montella, S., De Rosa, V. & La Cava, A. Leptin as a metabolic link to multiple sclerosis. Nat. Rev. Neurol. 6, 455–461 (2010).

27. Subramanian, A. et al. A Next Generation Connectivity Map: L1000 Platform and the First 1,000,000 Profiles. Cell 171, 1437–1452.e17 (2017).

28. Tissing, W. J. E. et al. Genomewide identification of prednisolone-responsive genes in acute lymphoblastic leukemia cells. Blood 109, 3929–3935 (2007).

29. Hu, Y. et al. Development of a Molecular Signature to Monitor Pharmacodynamic Responses Mediated by In Vivo Administration of Glucocorticoids. Arthritis Rheumatol. (Hoboken, N.J.) 70, 1331–1342 (2018).

30. Paragliola, R. M., Papi, G., Pontecorvi, A. & Corsello, S. M. Treatment with Synthetic Glucocorticoids and the Hypothalamus-Pituitary-Adrenal Axis. Int. J. Mol. Sci. 18, (2017).

31. Bali, U., Phillips, T., Hunt, H. & Unitt, J. FKBP5 mRNA Expression Is a Biomarker for GR Antagonism. J. Clin. Endocrinol. Metab. 101, 4305–4312 (2016).

32. Ysrraelit, M. C., Gaitán, M. I., Lopez, A. S. & Correale, J. Impaired hypothalamic-pituitary-adrenal axis activity in patients with multiple sclerosis. Neurology 71, 1948–1954 (2008).

33. Van Winsen, L. M. L. et al. Sensitivity to glucocorticoids is decreased in relapsing remitting multiple sclerosis. J. Clin. Endocrinol. Metab. 90, 734–740 (2005).

34. Wüst, S. et al. Peripheral T cells are the therapeutic targets of glucocorticoids in experimental autoimmune encephalomyelitis. J. Immunol. 180, 8434–8443 (2008).

35. Alcina, A. et al. Multiple sclerosis risk variant HLA-DRB1*1501 associates with high expression of DRB1 gene in different human populations. PLoS One 7, (2012).

36. Stefan, M. et al. Genetic-epigenetic dysregulation of thymic TSH receptor gene expression triggers thyroid autoimmunity. Proc. Natl. Acad. Sci. U. S. A. 111, 12562–12567 (2014).

37. Singh, S. P. et al. PLZF regulates CCR6 and is critical for the acquisition and maintenance of the Th17 phenotype in human cells. J. Immunol. 194, 4350–4361 (2015).

38. Ambele, M. A., Dessels, C., Durandt, C. & Pepper, M. S. Genome-wide analysis of gene expression during adipogenesis in human adipose-derived stromal cells reveals novel patterns of gene expression during adipocyte differentiation. Stem Cell Res. 16, 725–734 (2016).

39. Cepok, S. et al. Short-lived plasma blasts are the main B cell effector subset during the course of multiple sclerosis. Brain 128, 1667–1676 (2005).

40. Roostaei, T. et al. Defining the architecture of cerebrospinal fluid cellular communities in neuroinflammatory diseases. bioRxiv 2021.11.01.466797 (2021) doi:10.1101/2021.11.01.466797.

41. DeRijk, R. H., Eskandari, F. & Sternberg, E. M. Corticosteroid resistance in a subpopulation of multiple sclerosis patients as measured by ex vivo dexamethasone inhibition of LPS induced IL-6 production. J. Neuroimmunol. 151, 180–188 (2004).

42. Trombetta, J. J. et al. Preparation of Single-Cell RNA-Seq Libraries for Next Generation Sequencing. Curr. Protoc. Mol. Biol. 107, 4.22.1–4.22.17 (2014).

43. GTEx Consortium, K. G. et al. Human genomics. The Genotype-Tissue Expression (GTEx) pilot analysis: multitissue gene regulation in humans. Science 348, 648–60 (2015).

44. Dobin, A. et al. STAR: ultrafast universal RNA-seq aligner. Bioinformatics 29, 15–21 (2013).

45. DeLuca, D. S. et al. RNA-SeQC: RNA-seq metrics for quality control and process optimization. Bioinformatics 28, 1530–2 (2012).

46. Li, B. & Dewey, C. N. RSEM: accurate transcript quantification from RNA-Seq data with or without a reference genome. BMC Bioinformatics 12, 323 (2011).

47. Chang, C. C. et al. Second-generation PLINK: rising to the challenge of larger and richer datasets. Gigascience 4, 7 (2015).

48. Das, S. et al. Next-generation genotype imputation service and methods. Nat. Genet. 48, 1284–1287 (2016).

49. Jia, X. et al. Imputing Amino Acid Polymorphisms in Human Leukocyte Antigens. PLoS One 8, e64683 (2013).

50. Roostaei, T. et al. Proximal and distal effects of genetic susceptibility to multiple sclerosis on the T cell epigenome. Nat. Commun. 2021 121 12, 1–12 (2021).

51. Wu, Y. et al. Automated segmentation of multiple sclerosis lesion subtypes with multichannel MRI. Neuroimage 32, 1205–1215 (2006).

52. Ritchie, M. E. et al. limma powers differential expression analyses for RNA-sequencing and microarray studies. Nucleic Acids Res. 43, e47 (2015).

53. Watanabe, K., Taskesen, E., van Bochoven, A. & Posthuma, D. Functional mapping and annotation of genetic associations with FUMA. Nat. Commun. 8, 1826 (2017).

54. Chen, S., Zhou, Y., Chen, Y. & Gu, J. fastp: an ultra-fast all-in-one FASTQ preprocessor. Bioinformatics 34, i884–i890 (2018).

